# The genome of *Salmacisia buchloëana*, the parasitic puppetmaster pulling strings of sexual phenotypic monstrosities in buffalograss

**DOI:** 10.1101/2023.05.24.542141

**Authors:** Christopher W. Benson, Matthew R. Sheltra, David R. Huff

**Affiliations:** Department of Plant Science, Pennsylvania State University, University Park, PA, USA; Intercollegiate Graduate Degree Program in Plant Biology, Pennsylvania State University, University Park, PA, USA

**Keywords:** Extended phenotype, *Tilletia*, *Bouteloua dactyloides*, smut fungi, fungal pathogen, host manipulation

## Abstract

To complete its parasitic lifecycle, *Salmacisia buchloëana*, a biotrophic fungus, manipulates reproductive organ development, meristem determinacy, and resource allocation in its dioecious plant host, buffalograss (*Bouteloua dactyloides;* Poaceae). To gain insight into *S. buchloëana’s* ability to manipulate its host, we sequenced and assembled the 20.1 Mb genome of *S. buchloëana* into 22 chromosome-level pseudomolecules. Phylogenetic analysis suggests that *S. buchloëana* is nested within the genus *Tilletia* and diverged from *T. caries* and *T. walkeri ∼*40 million years ago. We find that *S. buchloëana* has a novel chromosome arm with no syntenic relationship to other publicly available *Tilletia* genomes and that genes on the novel arm are upregulated upon infection, suggesting that this unique chromosomal segment may have played a critical role in *S. buchloëana’s* evolution and host specificity. *Salmacisia buchloëana* has one of the largest fractions of serine peptidases (1.53% of the proteome) and one of the highest GC contents (62.3%) in all classified fungi. Analysis of codon base composition indicated that GC content is controlled more by selective constraints than directional mutation and that *S. buchloëana* has a unique bias for the serine codon UCG. Finally, we identify three inteins within the *S. buchloëana* genome, two of which are located in a gene often used in fungal taxonomy. The genomic and transcriptomic resources generated here will aid plant pathologists and breeders by providing insight into the extracellular components contributing to sex determination in dioecious grasses.

## Introduction

*Salmacisia buchloëana* Huff & Chandra (syn. *Tilletia buchloëana* Kellerman & Swingle) is a fungal biotroph that spends most of its lifecycle growing intercellularly in its plant host, buffalograss (*Bouteloua dactyloides* [Nutt.] Columbus; syn. *Buchloë dactyloides* [Nutt.] Engelmann). *Salmacisia buchloëana* completes its lifecycle by producing teliospores in buffalograss ovaries, but because buffalograss is dioecious, the reproductive capacity of the fungus is restricted to only those plants with female floral anatomy (*i.e.*, half of the host population). To mitigate this reproductive bottleneck, *S. buchloëana* has evolved to induce female floral organs (stigmas, styles, ovaries) in the flowers of genetically male buffalograss for the purpose of teliospore production and ultimately completion of its lifecycle (Chandra and Huff, 2008). In this way, *S. buchloëana* hijacks the genetic machinery involved with floral development in its grass host to further its own reproductive potential (Fig. 1).

**Fig. 1.**
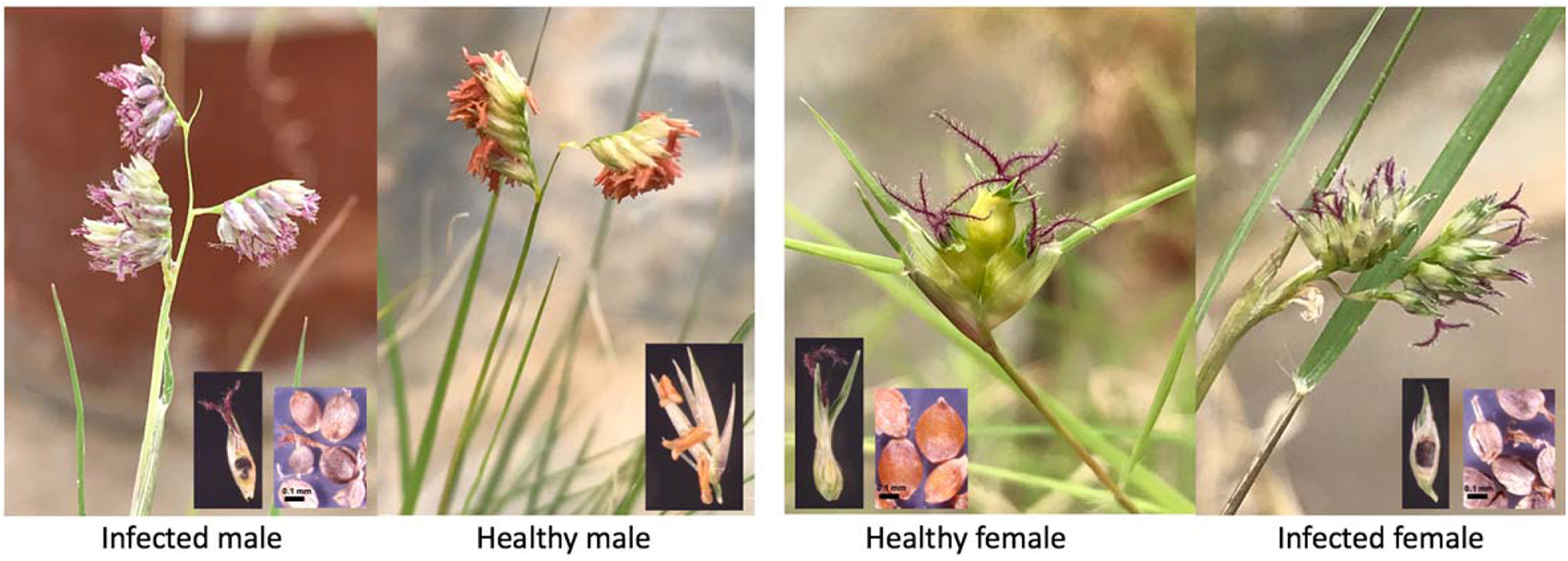
Dioecious buffalograss either infected with *Salmacisia buchloëana* or healthy (mock-infected). Infection with *S. buchloëana* induces the development of female floral organs (pistils) in the flowers of male plants. Pistils and stamens are easily visible with their purple feathery stigmas and orange anthers, respectively. The inset images in the bottom corners show buffalograss florets and ovaries. Healthy males do not produce ovaries, so are not depicted. The fungal-induced ovaries of infected plants are filled with teliospores and mature into ‘bunt balls’ that are, on average, smaller than seed from uninfected female ovaries (scale bar = 0.1 mm).

Dioecious buffalograss with unisexual floral arrangement likely evolved from a hermaphroditic ancestor with bisexual flowers (Kinney *et al*., 2007). As a result, unisexual buffalograss flowers contain nonfunctional rudiments of the opposite floral organ (*i.e.*, vestigial stamens in female plants and pistil primordia in male plants; Chandra and Huff, 2010). Infection with *S. buchloëana* overrides buffalograss’ unisexual reproductive biology to induce the development of the otherwise aborted floral organs, resulting in a bisexual flower (Chandra and Huff, 2008). The induced ovaries of male plants are easily visible and play an important role in *S. buchloëana’s* reproductive lifecycle, but the induced stamens of female plants are underdeveloped and are not involved in sporulation, suggesting that they may be an off-target byproduct of fungal manipulation (Chandra and Huff, 2010). In addition to manipulating floral architecture, Chandra and Huff (2014), found that *S. buchloëana* influences broad physiological traits in its host, including resource partitioning and meristem determinacy with infected plants having increased sexual allocation at the expense of vegetative allocation. It is unclear if multiple buffalograss traits are specifically targeted by *S. buchloëana* (*i.e.,* multidimensional phenotypes; Thomas *et al*., 2010; van Houte, Ros, and van Oers 2013; Poulin 2013; Cézilly *et al*., 2013), or if the fungus manipulates a single trait and other phenotypes are incidental costs of manipulation. In either case, the altered phenotypes of buffalograss are the result of manipulation by *S. buchloëana* and therefore represent the ‘extended phenotype’ of *S. buchloëana* (Vyas, 2015; Dawkins, 2016; Henry *et al*., 2021).

*Salmacisia* is a monotypic genus and falls within the order Tilletiales (Basidiomycota, Ustilaginomycotina, Exobasidiomycetes) that includes ca. 191 species of fungi, many of which produce teliospores in the ovaries of their grass (Poaceae) hosts (He *et al*., 2019). Species in the Tilletiales are characterized by forming dark pigmented spores with a pungent odor and commonly referred to as ‘smut fungi’. To our knowledge, *S. buchloëana* is the only species within the Tilletiales known to infect a dioecious host and thereby, the only Tilletiales to induce ovaries in male plants. Infection with *S. buchloëana* is uncommon in nature but has been reported throughout the southern Great Plains of the United States and central Mexico (Huff *et al*., 1987).

Here, we compare the genomic features of fungi in the Tilletiales to identify novel components of the *S. buchloëana* genome that might play a role in its unique ability to manipulate host sex organ identity and other extended phenotypes. The findings and genomic resources presented here will guide further analyses into the fine-tuned regulatory pathways associated with sex manipulation in the *Salmacisia*-buffalograss pathosystem.

## Results

### Genome assembly and annotation

The OK1 strain of *S. buchloëana* was sequenced to 51× coverage using the PacBio Sequel system. The Canu genome assembly (Koren *et al*., 2017) resulted in 30 contigs, two of which were circular and six were singletons (represented by one sequence). One of the circular contigs was identified as the complete 86,026 bp mitochondrial genome (Supplementary Fig. 1) and the other aligned to a PacBio internal control and was subsequently removed. The six singletons were independently aligned to the 22 remaining contigs to check for their representation in the consensus contigs. All six singletons shared >97% sequence identity to the consensus contigs and were removed from the assembly. The 22 remaining contigs ranged between 0.54 to 1.46 Mb in length (Supplementary Table 1) and were similar in size and structure to the full-length chromosomes of model fungi, *Ustilago maydis* (Kämper *et al*., 2006) and *U. bromivora* (Rabe *et al*., 2006).

We used the Benchmarking Universal Single-Copy Orthologs (BUSCO) software to scan for conserved fungal genes and found that the genome contained 95.8% of the 1,335 single-copy orthologs in the Basidiomycota, suggesting that the 22 scaffolds represent the chromosome-level pseudomolecules of *S. buchloëana* (Table 1). In addition, we scanned for telomeric repeat sequences at the ends of *S. buchloëana* pseudomolecules to further validate the chromosome-level assembly. Plant, mammal, and fungal chromosomes typically end in (TTAGGG)n repeats (Meyne *et al*., 1990; Wu *et al*., 2010). We found that 18 of the 22 *S. buchloëana* chromosomes contained canonical (TTAGGG)n-3’ telomeric repeat sequences at both ends while the remaining 4 pseudomolecules possessed telomeric repeats at one end, further suggesting that the genome assembly spans the near full length of *S. buchloëana’s* chromosomes (Supplementary Table 1).

**Table 1.** Genomic features of *Salmacisia buchloëana* compared to related fungal genomes.

| Genomes:<br>NCBI WGS ID | *S. buchloëana*<br>MOEQ01 | *T. horrida*<br>LAXH01 | *T. caries*<br>LWDD01 | *T. controversa*<br>LWDE01 | *T. laevis*<br>RDSF01 | *T. indica*<br>LWDF01 | *T. walkeri*<br>LWDG01 | *U. maydis*<br>AACP02 |
| --- | --- | --- | --- | --- | --- | --- | --- | --- |
| Size (Mb) | 20.1 | 20.1 | 29.5 | 28.8 | 28.8 | 30.4 | 23.3 | 19.7 |
| Scaffold No. | 22 | 767 | 2,888 | 3,586 | 3,961 | 1,666 | 972 | 27 |
| Scaffold N50 (bp) | 901,006 | 75,652 | 32,675 | 14,841 | 13,920 | 83,419 | 59,453 | 884,984 |
| BUSCO of genome | 93.6 | 93.9 | 93.1 | 92.4 | 92.5 | 93.4 | 94.6 | 99.3 |
| % GC | 62.3 | 55.8 | 56.4 | 56.9 | 56.6 | 54.5 | 54.9 | 54.0 |
| Protein coding genes | 6,379 | 6,108 | 10,204 | 9,860 | 9,799 | 9,548 | 7,970 | 6,782 |
| BUSCO of protein annotation | 95.8 | 88.7 | 96.6 | 95.9 | 96.3 | 97.1 | 97.4 | 99.6 |
| tRNAs | 56 | 77 | 78 | 80 | 102 | 68 | 65 | 102 |
| Gene density (#/Mb) | 317 | 303 | 346 | 342 | 340 | 314 | 342 | 344 |
| Unique proteins | 949 | 77 | 1,748 | 1,472 | 731 | 2,072 | 1,030 | 2,313 |
| % SSRs† | 3.3 | 2.3 | 2.8 | 2.9 | 2.5 | 2.3 | 2.4 | 1.9 |
| Avg. transcript length | 1,886 | 1,705 | 1,579 | 1,596 | 1,575 | 1,711 | 1,724 | 1,744 |
| % genome transcribed | 60 | 52 | 56 | 57 | 54 | 57 | 55 | 61 |
| Secreted proteins‡ | 852 | 861 | 1457 | 1413 | 1378 | 1346 | 1144 | 1009 |
| GPI anchor proteins | 16 | 22 | 18 | 21 | 21 | 16 | 18 | 17 |
| Secretory proteins<br>(% proteome) | 836<br>(13.1) | 839<br>(13.7) | 1439<br>(14.1) | 1392<br>(14.1) | 1357<br>(13.8) | 1330<br>(13.9) | 1126<br>(14.1) | 992<br>(14.6) |
| Effector proteins<br>(% proteome) | 256<br>(4.0) | 285<br>(4.66) | 467<br>(4.6) | 446<br>(4.5) | 429<br>(4.37) | 419<br>(4.4) | 331<br>(4.2) | 343<br>(5.1) |

The *S. buchloëana* genome is 5.7% repetitive DNA, with the largest repeat categories being simple sequence repeats (3.3%) and Long Terminal Repeat (LTR) retrotransposons (1.9%; Supplementary Table 2). Proteins were predicted using *S. buchloëana* transcriptomic sequences to guide *ab initio* gene prediction (Cantarel *et al*., 2008). Genome annotation showed that, relative to related fungi (Table 1), *S. buchloëana* has the fewest predicted protein coding genes (6,379), fewest predicted tRNAs (56), fewest unique proteins (949), and fewest number of predicted secreted proteins and effectors at 836 (13.1% of the proteome) and 256 (4.0% of the proteome), respectively. However, *S. buchloëana* has the longest average transcript length and a relatively high percent of the genome transcribed. Interestingly, we find that *S. buchloëana* has retained genes in the sulfur and nitrogen metabolic pathways that are typically missing in obligate biotrophic fungi (Sharma *et al*., 2015; Jiang *et al*., 2013), indicating that *S. buchloëana* may survive outside its host in certain environmental conditions (Supplementary Fig. 2).

Centromeric sequences typically have lower GC content (Diner *et al*., 2017), lower gene density, and are enriched with long tandem repeats (Melters *et al*., 2013). We scanned for these three features across *S. buchloëana* chromosomes to identify putative centromeric regions (Fig. 2; Supplementary Fig. 3). The size of the 22 predicted centromeric regions ranged from 32 to 181 kb in length. This proposed range of centromere sizes is in agreement with other fungal centromeres measured using a specialized histone H3 variant, CENP-A (the acid test for centromeric locations; Smith, 2002). All predicted *S. buchloëana* centromeres were some version of metacentric or acrocentric with the exception of chromosomes 17 and 22, which were telocentric.

**Fig. 2.**
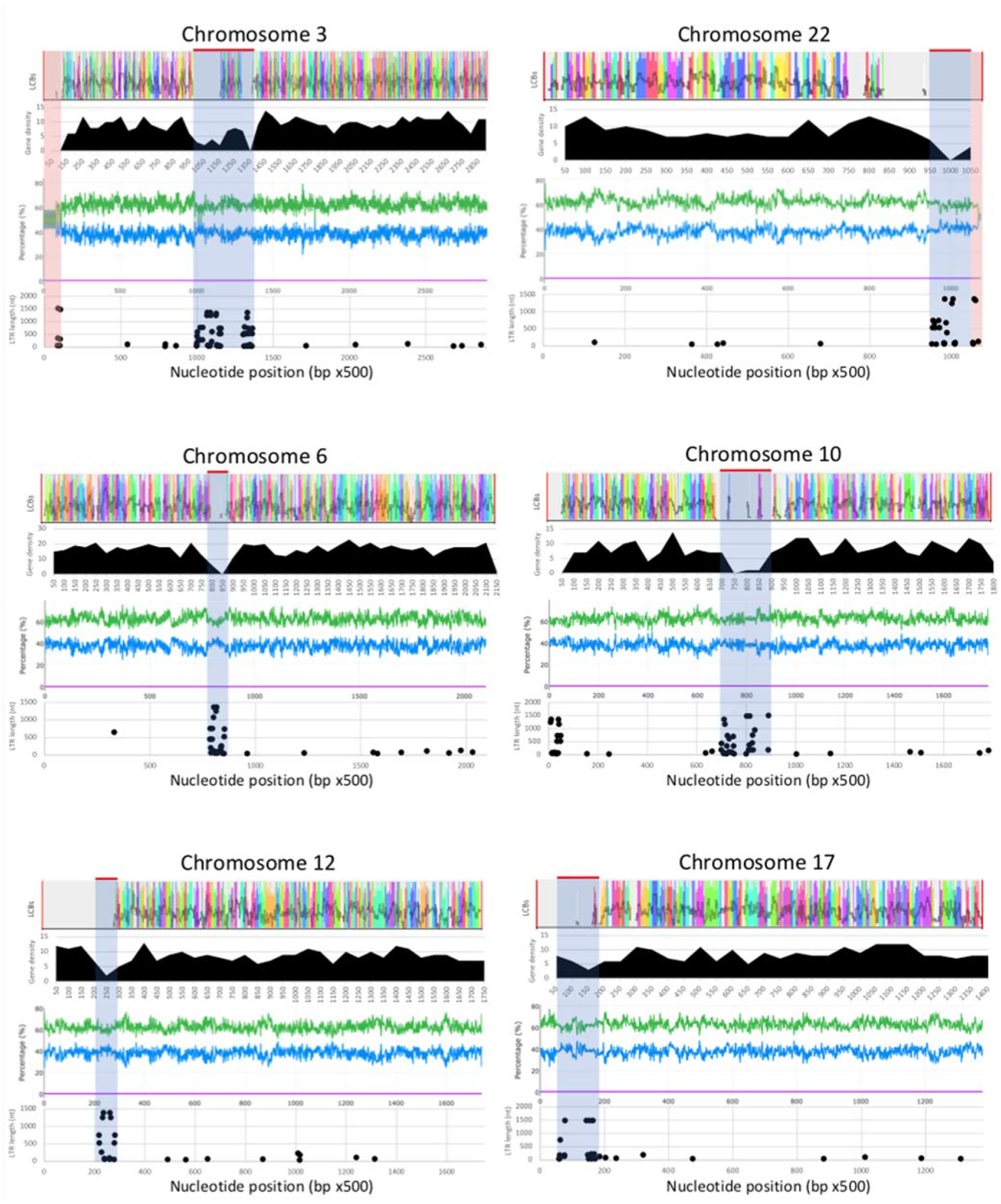
Genome features across six of the 22 chromosomes of *Salmacisia buchloëana*. For each chromosome, from top to bottom, graphs show (1) The Local Colinear Blocks (LCBs) of *S. buchloëana* sequences compared to five *Tilletia* genomes (see methods), where colors represent shared synteny, (2) Gene density across a 25 kb sliding window (black histogram), (3) Percent GC (green) and AT content (blue) per 500 nucleotides (center graph), and (4) LTR retrotransposon location and length (dot plot). Putative centromeric regions are indicated with a gray shaded box, while the shaded red boxes highlight the two ribosomal DNA sequences with reduced GC content.

### Phylogenetic Analysis and Molecular Dating

*Salmacisia buchloëana* shares similar morphological characteristics with species in the genus *Tilletia* and was initially placed within *Tilletia* (Kellerman and Swingle, 1889) but later reclassified and renamed based on host taxonomy, spore ornamentation, and DNA sequence analysis (Chandra and Huff, 2008). Piątek *et al*., (2016) conducted a phylogenetic analysis using 28S ribosomal DNA (rDNA) sequences and also found that *S. buchloëana* resides phylogenetically outside of the *Tilletia*, while Jayawardena *et al*., (2019) conducted a similar study and found that *S. buchloëana* placed within the *Tilletia* genus. The discrepancy in phylogenetic placement may not be surprising since, in all three studies, *S. buchloëana* resides on a long phylogenetic branch, indicating a high level of molecular divergence from its nearest ancestors, raising the possibility of incorrect placement due to error associated with ‘long branch attraction’ (Felsenstein, 1978). Here, we used 328 pairs of single-copy orthologous genes spanning the Ustilaginomycotina and found that *S. buchloëana* fell within the genus *Tilletia* (Fig. 3).

**Fig. 3.**
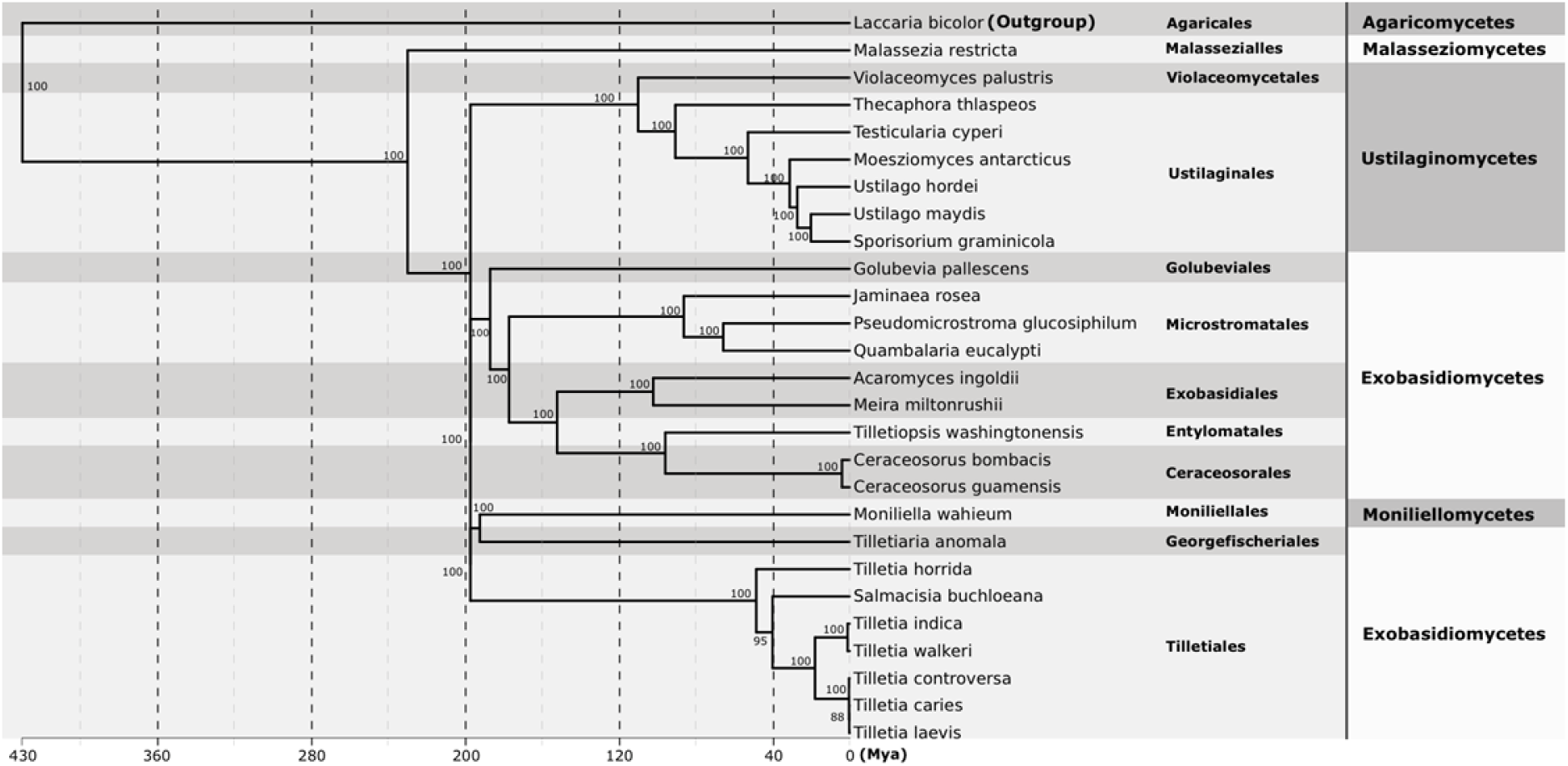
A phylogenetic classification of fungal genomes in the Ustilaginomycotina based on 328 single-copy orthologs. The tree was generated using RAxML with 100 bootstraps. Bootstrap probabilities are shown above branches. Branch lengths are scaled to the divergent time estimates.

The common ancestor of *S. buchloëana* and outgroup *Laccaria bicolor* is estimated to have diverged early in the evolution of the Basidiomycota, ∼430 Million years ago (Mya; Zhao *et al*., 2017; He *et al*., 2019). We used the divergence of *S. buchloëana* and *L. bicolor* to calibrate our molecular dating and found that *S. buchloëana* diverged from *T. horrida* 48 Mya and diverged from the other five species of *Tilletia* (*T. indica, T. walkeri, T. caries, T. controversa,* and *T. laevis*) 40 million years ago. Our analysis suggests that the *T. caries* and *T. walkeri* clades diverged from each other 18 Mya.

### Analysis of high GC content in *Salmacisia buchloëana*

GC content can range from 13 to 80% in bacteria but is typically less than 50% for plants, animals, and fungi (Li and Du, 2014). Most coding regions have a higher GC content than non-coding regions and for this reason, many researchers have investigated the cause and utility of GC content variation. Chromosomal regions with high GC content have been termed ‘isochores’ in animals and are described as giving stability and structure to the genome (Vinogradov, 2003). GC content has been implicated in molecular phenomena, including GC-biased gene conversion (Long *et al*., 2018), reduced DNA denaturation in GC-rich regions (Fryxell *et al*., 2000), and the negative relationship between GC content and mutation (Wolfe *et al*., 1989).

Species in the Basidiomycota have the highest GC content among fungi (mean = 54.6%; Storck, 1966). *Salmacisia buchloëana* (GC% = 62.3 whole genome; 63.7 genic sequences) has the highest GC content compared to any of its closest relatives and was among the highest in the Basidiomycota; less than *Anthracocystis flocculosa* (65.3%) but higher than *Sporobolomyces salmonicolor* (61.3%) and *Rhodotorula mucilaginosa* (59.9%). The distribution of GC content is uniformly high along *S. buchloëana’s* 22 chromosomes, with minor exceptions. The large and small rDNA subunits on chromosomes 3 and 22 represented the regions with the highest AT contents across the entire 20.1 Mb genome (Fig. 2; Supplementary Fig. 3). We compared the distribution of GC content of other fungi in the Basidiomycota and found that reduced GC content is maintained in the rDNA sequence of each of the species that we analyzed (Supplementary Fig. 4), suggesting that rDNA is resistant to variation in GC content in the Basidiomycota and may be under purifying selection to maintain this pattern of oscillating GC content at about a 50% level. We do not detect a strong signal of repeat-induced point mutations (RIP) in the *S. buchloëana* genome (0.03%), a result that was expected since RIP regions typically have higher AT content (Selker and Stevens, 1985; Hane and Oliver, 2008).

### GC Content and Codon Usage

We analyzed an orthologous gene set from five of the six *Tilletia* species that included at least 948 of 985 orthologous genes depending on the species (Note: the *T. laevis* genome was not publicly available at the time these analyses were performed). Using this orthologous gene set, we performed a neutrality test to compare the GC content of the 1^st^ and 2^nd^ codon positions (GC12) against that of the 3^rd^ position (GC3; *i.e.,* the synonymous codon position; Fig. 4). We found that *S. buchloëana* had the highest GC content for coding, genome, or non-coding sequences (Fig. 4A) and the highest relative selective constraints on its GC content (Fig. 4B) compared to any of the other five *Tilletia* species. While the level of selective constraints seemed to be correlated with overall GC content, there was an exception with *T. horrida* in that it showed a relatively low GC content but a relatively high selective constraint value (Fig. 4A vs B). Given the assumption that mutation rates are in equilibrium, one possible explanation is that selective constraints are keeping the GC content of *T. horrida* lower than its close relatives.

**Fig. 4.**
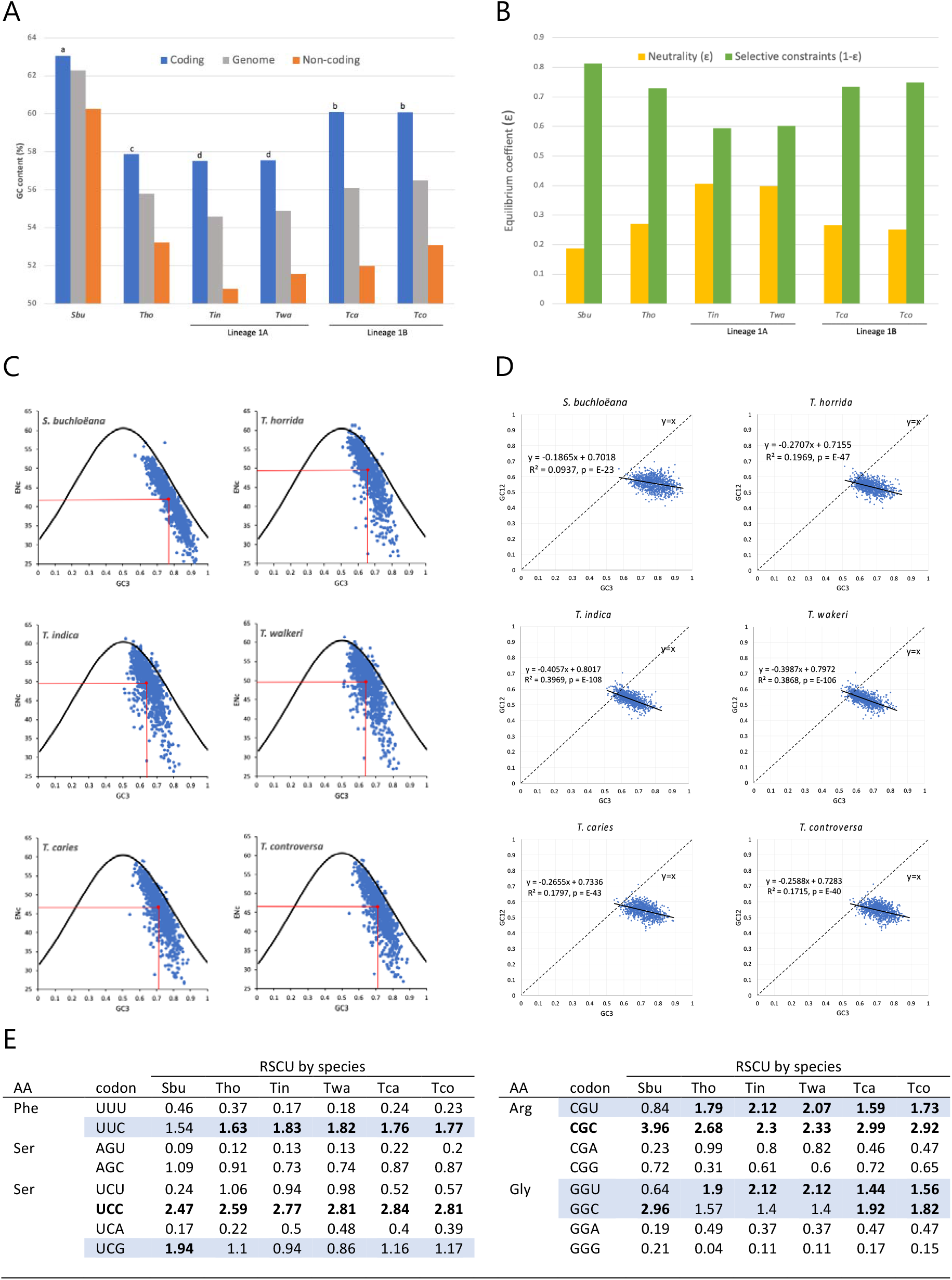
GC content and codon usage in *Salmacisia buchloëana*. **(A)** Percent GC content in *S. buchloëana* and other species in the genus *Tilletia*. GC content (%) of coding (orthologous gene set), non-coding and genomic sequence. Coding means topped by different letters are significantly different at the p=0.0001 level of significance. (**B)** Neutrality vs selection of codon GC12 content. Evolutionary modeling of GC content at the 1^st^ and 2^nd^ codon position (GC12) of the orthologous gene set. The absolute value of the mutation-selection equilibrium coefficient ɛ (approximated by the slope of the neutrality plots) equals the relative effects of neutrality while 1 – ɛ equals the relative effect of selection constraints governing GC12 content. (**C)** Effective number of codons (ENc) vs GC content at the 3^rd^ codon position (GC3) of the orthologous gene set coding sequences (blue dots). Black line represents the theoretical limit of ENc, red lines indicate mean values of ENc and GC3. (**D)** Neutrality plots (GC12 vs GC3) of the orthologous gene set coding sequences (blue dots). Linear regression (solid black line) equation and coefficient of determination (R^2^) indicated. Theoretically complete equilibrium with directional mutation (y=x) is represented by dashed line. (**E)** Relative synonymous codon usage (RSCU) for the orthologous gene set from *Salmacisia buchloëana* and the five *Tilletia* species. RSCU values in bold are significantly (p<0.01) biased as determined by a two-way Chi-square contingence test in CodonW. Codons in bold are significantly biased across all species. Highlighted codons denote differences among species (see Supplementary Table 3 for the full RSCU table).

The effective number of codons (ENc; Fig. 4C) and the neutrality plots (GC12 vs GC3; Fig. 4D) suggest that *S. buchloëana* has less codon bias than the other *Tilletias* (Fig. 4C). The five *Tilletia* species appeared to have more genes that were further in distance from the ENc equilibrium line (Fig. 4C) and had higher mean ENc and steeper regression slopes (Fig. 4D) than *S. buchloëana*, suggesting that they have more codon bias. Similarly, analysis of the relative synonymous codon usage (RSCU) found more codon bias for the five *Tilletia* species than for *S. buchloëana*. Out of a total of 21 biased codons, 20 were biased in *T. caries* and *T. controversa*, 19 were biased in *T. horrida*, *T. indica*, and *T. walkeri*, and 18 codons were biased in *S. buchloëana* (Supplementary Table 3). Sixteen (76%) of the 21 total codon biases observed were shared across all six species. The five instances of biased discrepancy among species all involve *S. buchloëana* (Fig. 4E). In three of the five instances, *S. buchloëana* lacked a significant codon bias that all other *Tilletia* species shared (UUC, Phe; CGU, Arg; GGU, Gly). In one instance, *S. buchloëana* shared a bias (GGC, Gly) with the other two systemically infecting fungi, *T. caries* and *T. controversa*. The fifth and final codon bias was only present in *S. buchloëana*, *i.e*., the UCG codon for serine.

### CAZymes and MEROPS

Carbohydrate-Activated enZymes (CAZymes) are involved with the synthesis and degradation of polysaccharides and glycoconjugates (Park *et al*., 2010). Biotrophic fungi such as *S. buchloëana* and the *Tilletias* rely on their hosts for survival and completion of their fungal lifecycle, and therefore typically have fewer CAZymes than hemibiotrophic, saprotrophic, and necrotrophic fungi. Among the fungal biotrophs in this study, *S. buchloëana* had an intermediate number of CAZymes at 313 with modest depletions in all enzyme classes except the largest class, glycoside hydrolases (GHs; Supplementary Fig. 5A and B). The CAZyme profile of *S. buchloëana* is similar to the *Tilletias*, likely due to their shared evolution and biotrophic relationship to their hosts.

Fungal peptidases (proteases) are necessary for digestion of protein substrates and are often secreted into the environment for the breakdown of external protein targets. Secreted peptidases are essential for pathogenicity and considered virulent to the plant host. Classification of peptidases and their inhibitors are available at the MEROPS database (Rawlings *et al*., 2018).

The distribution of peptidase families in *S. buchloëana* is similar to the *Tilletias*, with only a couple noteworthy exceptions, one being the presence of inteins (N09s). Genomes are known to contain selfish genetic elements that promote their own replication at the expense of the host, including transposable elements, self-promoting plasmids, and B chromosomes (Werren, 2011). Inteins (*<u>int</u>*ervening prot*<u>eins</u>*) are a special class of selfish genetic elements and similar in concept to the introns of DNA (Shah and Muira, 2014). Inteins range in size from 134 to 1,065 amino acids and are mostly found in bacteria and archaea (Green *et al*., 2018). Currently, there are 257 known inteins that have been identified in 231 species of eukaryotes, with 15 inteins being found in the Basidiomycota, primarily in the human pathogen *Cryptococcus* spp. and the bunt genus *Tilletia* (Green *et al*., 2018). Thus, it is uncommon for eukaryotic species to possess an intein and rare to contain more than one intein.

A total of three genes within the genome of *S. buchloëana* were found to contain inteins, namely in the pre-mRNA-splicing process factor 8 (Prp8; MOEQ 005882) gene on chromosome 8, the DNA-dependent RNA polymerase 2 (RPB2; MOEQ 004009) gene on chromosome 3, and the DNA-dependent RNA polymerase 2-like (RPB2-like; MOEQ 002009) gene on chromosome 15. The amino acid sequences of these three inteins will be referred to as SbuPrp8i, SbuRPB2i, and SbuRPB2-likei, respectively. Inteins are transmitted to their hosts both vertically and horizontally (Green *et al*., 2018). Thus, phylogenic trees do not necessarily represent an accurate portrait of the relatedness among inteins across different fungal species. However, the maximum likelihood phylogenetic trees representing the SbuPrp8i intein or SbuRPB2i and SbuRPB2-likei inteins in fungi did cluster according to phyla (Fig. 5A and B). Like all inteins, the SbuPrp8i, SbuRPB2i, and SbuRPB2-likei begin with a cystine residue (C-1) and end with an asparagine residue (N-284, N-397, and N-423, respectively) (Fig. 5C and E). SbuPrp8i also contains a LAGLIDADG-type homing endonuclease domain as well as an N-splicing domain (Blocks A&B) and a C-splicing domain (Blocks F&G) (Fig. 5C). According to Duan *et al*. (1997), blocks C and E are the original LAGLIDADG motifs and each contains an endonuclease active site Asp (D) or Glu (E) while block D contains a putative active site Lys (K). However, as a result of mutations, several inteins from other species examined were found to contain only partial motifs. The Prp8 inteins of *Cryptococcus gatti* and *C. neoformans* completely lack a LAGLIDADG-type homing endonuclease and, as such, are referred to as ‘mini-inteins’. Interestingly, the two inteins, RPB2i and RPB2-likei, identified here (Fig. 5E) reside between two frequently used reverse primers (bRPB2-7R and bRPB2-7.1R) for the amplification of the fungal RPB2 gene which is commonly used for phylogenetic analysis of fungi (Sun *et al*., 2009). Thus, the presence of either RPB2i or RPB2-likei has the potential to alter RPB2 amplicon sequence information and hence alter the phylogenetic placement of species containing these specific inteins. Reverse primer bRPB2-7R is located in the RPB2 N-extein typically six residues from RPB2 intein block A, while reverse primer bRPB2-7.1R is located typically one residue from RPB2 intein block G in the RPB2 C-extein. Fungal taxonomists should make note of this finding for their future use of the RPB2 gene in phylogenetic analyses.

**Fig. 5.**
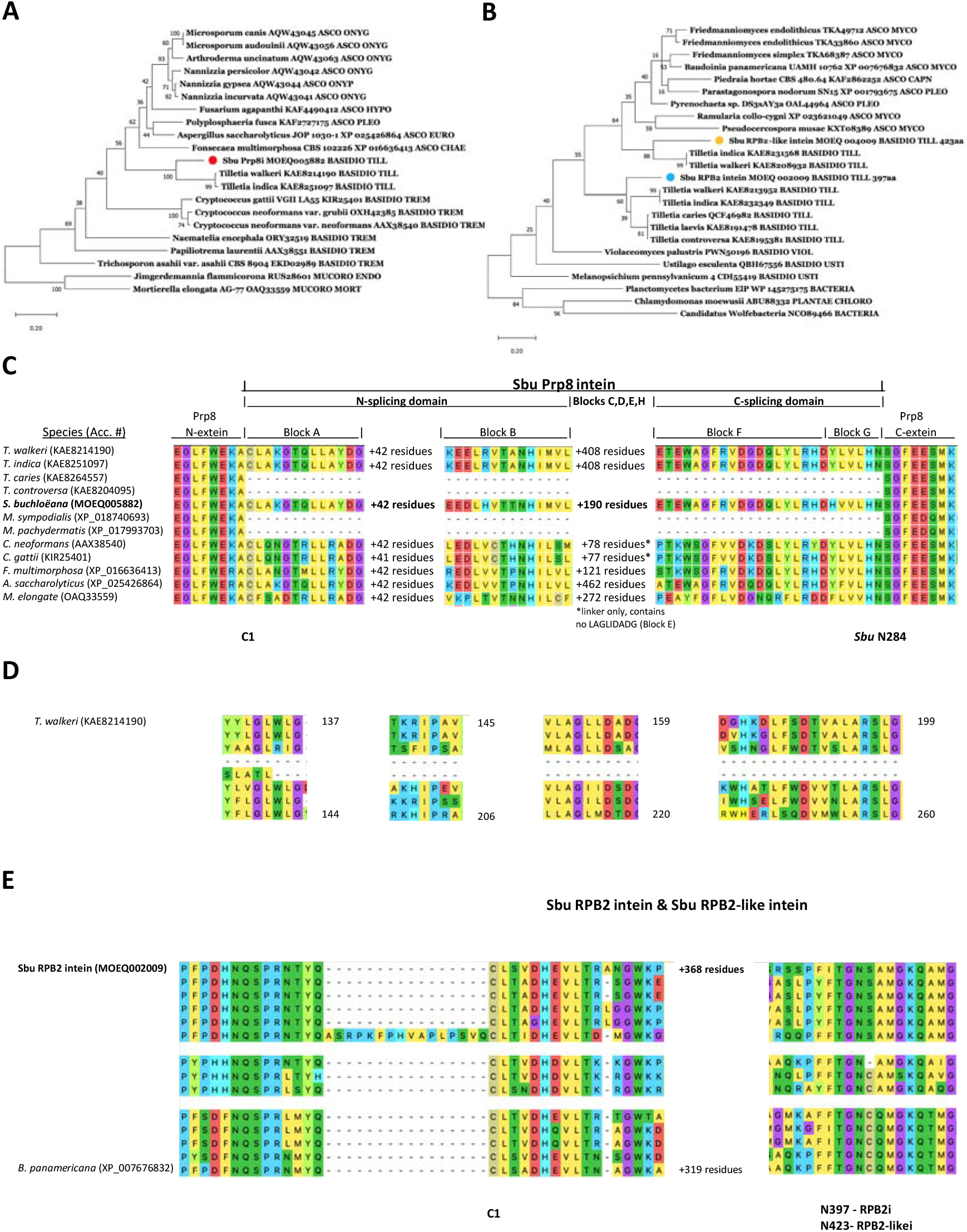
Inteins of *Salmacisia buchloëana* include Prp8i, RPB2i and RPB2-likei. **(A)** Phylogenetic tree of the *S. buchloëana* Prp8 intein (red dot) with selected inteins from other fungi (highest log likelihood tree, -2991.50; 550 bootstraps with bs >50 indicated; phylum and order of each fungal host is abbreviated; Abbreviations: ASCO, Ascomycota [CHAE, Chaetothyriales; EURO, Eurotiales; HYPO, Hypocreales; ONYG, Onygenales; PLEO, Pleosporales]; BASIDIO, Basdiomycota [TILL, Tilletiales; TREM, Tremellales; USTI, Ustilaginales]; MUCORO, Mucoromycota [ENDO, Endogonales; MORT, Mortierellales]). **(B)** Phylogenetic tree of the *Salmacisia* RPB2 intein (blue dot) and RPB2-like intein (brown dot) with selected inteins from fungi and bacteria (highest log likelihood tree, -1407.72; 550 bootstraps with bs >50 indicated; phylum and order of each host species is abbreviated; Abbreviations: ASCO, Ascomycota [CAPN, Capnodiales; MYCO, Mycocaliciales; PLEO, Pleosporales]; BASIDIO, Basdiomycota [TILL, Tilletiales; USTI, Ustilaginales]; PLANTAE CHLORO, Chlorodendrales). **(C)** Fungal species containing the Prp8 intein share certain domains including the N-splicing domain (Blocks A&B), the C-splicing domain (Blocks F&G) and variable amounts of linker between blocks B and F that may contain a LAGLIDADG-type homing endonuclease. The genomes of *Tilletia caries*, *T. controversa*, *Malassezia sympodialis and M. pachydermatis*, do not contain the Prp8 intein for comparison. **(D)** The DOD (LAGLIDADG) homing endonuclease helix motifs (Blocks C, D, E, H) of Prp8 inteins from selected fungal species. **(E)** RPB2 and RPB2-like inteins and exteins along with positions for two of the most frequently used reverse primers (bRPB2-7R and bRPB2-7.1R) for the amplification of the fungal RPB2 gene in phylogenetic studies. Reverse primer bRPB2-7R is in the RPB2 N-extein typically six residuals from RPB2 intein Block A, while reverse primer bRPB2-7.1R is located typically 1 residual from RPB2 intein Block G in the RPB2 C-extein.

Our analysis of peptidases also revealed that *S. buchloëana* has one of the largest fractions of serine peptidases in all classified fungi to date (1.53% of the proteome), falling just behind the highest recorded fungus, ascomycete *Torrubiella hemipterigena* at 1.56% (Muszewska *et al*., 2017). *Salmacisia buchloëana’s* enrichment in serine peptidases is primarily due to an abundance of genes in the subtilisin family (Supplementary Fig. 5C and D). Subtilisins (S08s) are involved with cellular degradation and hormone activation and are often secreted proteins found to be enriched in fungi with a pathogenic lifestyle (Muszewska *et al*., 2017; Leger *et al*., 1997). Interestingly, *S. buchloëana* has a noticeably higher number of predicted subtilisins than any of the closely related species that we examined (*S. buchloëana* = 23, *Tilletia* sp. ≤ 14). Many of *S. buchloëana’s* subtilisins are products of gene duplication and are unique to the species (Fig. 6E). The abundance of subtilisins in the *S. buchloëana* genome suggests that they may have played a functional role in its host specificity.

**Fig. 6.**
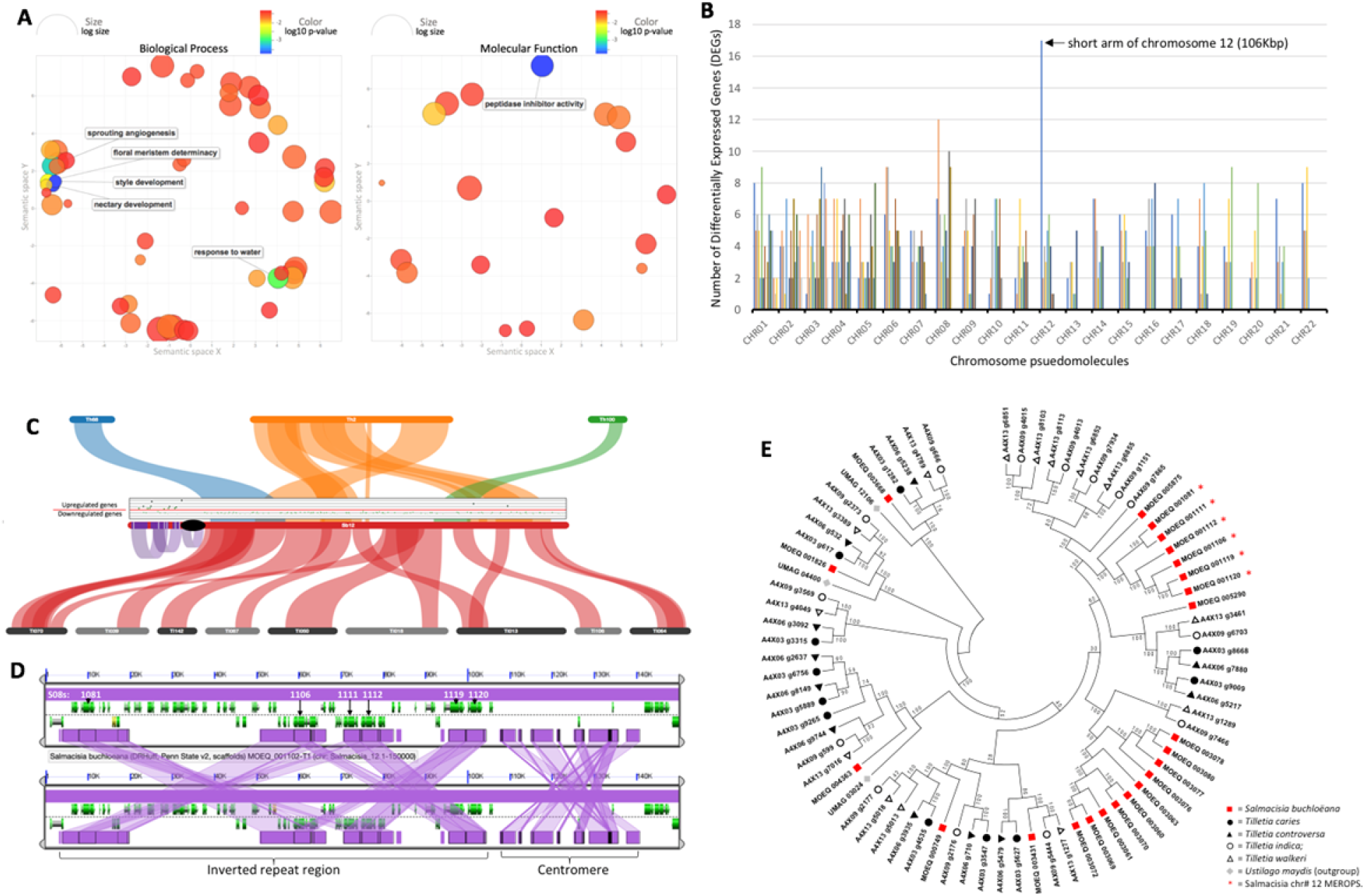
Features of host manipulation and the unique short arm of chromosome 12. **(A)** Analysis of gene ontologies (GOs) shows functionally enriched categories of male buffalograss when infected with the sex-altering fungus, *Salmacisia buchloëana.* Enriched GOs are clustered if their function is semantically similar. Only enrichments with a log10 p-value ≤ -2.5 are shown. On the left, GOs associated with biological process and on the right, GOs with a molecular function. Color of the bubbles indicates the p-value associated with the term, and size indicates the frequency of the GO term in the Gene Ontology Annotation (GOA) database (General GO terms have larger bubbles). **(B)** The distribution of up-regulated genes across the chromosomes of *S. buchloëana* (log_2_ fold change ≥1.5; false discovery rate ≤ 0.05) when the fungus is grown in its host rather than in culture (see methods). Chromosome distributions used a sliding, non-overlapping 100 kb window. (C) Gene expression (scatter plot) and syntenic comparisons (colored ribbons) across chromosome 12 highlights sequence and functional novelty in *S. buchloëana*. *Tilletia horrida* scaffolds (Th68, Th2, and Th100) with synteny to chromosome 12 (top) and *T. indica* (Ti018-Ti142) scaffolds with synteny to *S. buchloëana* (bottom) illustrate that the short arm of chromosome 12 is unique to *S. buchloëana*. Purple ribbons on *S. buchloëana* chromosome 12 show self-syntenic duplicated regions present as inverted repetitive sequence and the black ellipse is the putative centromeric sequence. White bands mark the location of subtilisin genes (S08). The scatterplot above *S. buchloëana* chromosome 12 depicts gene expression along the chromosome during buffalograss infection (log_2_ fold change ≥1.5; false discovery rate ≤ 0.05). Above the red line are genes that are upregulated during infection and below the line are genes downregulated during infection. (D) Expanded view of the duplicated and inverted repeats of S08s on the short arm of chromosome 12. (E) Evolutionary relationships of S08s among Tilletiales. Red squares = *S. buchloëana*; black circles = *T. caries*; black triangles = *T. controversa*; open circles = *T. indica*; open triangles = *T. walkeri*; grey diamonds = *Ustilago maydis*. Red asterisks are located on chromosome 12 and are unique to *S. buchloëana*.

### Sex alteration of the host

The ability for fungi to manipulate the sex of their hosts is rare among biotrophs but not entirely unique to *S. buchloëana*. The best described example is anther-smut (*Microbotryum* spp.), a genus of fungi that infects plants in the Caryophyllaceae family and replaces pollen with fungal spores in developing flowers (Hood *et al*., 2010; Kemler *et al*., 2020). In dioecious Caryophyllaceae (ex. *Silene latifolia* and *S. dioica*), infection with *Microbotryum* spp. causes anthers to develop in genetically female plants (Uchida *et al*., 2003). Genomic and transcriptomic surveys have helped identify cell-wall degrading enzymes, secondary lipases, glycosyltransferases, and other enzymes that might play a role in *Microbotryum’s* biotrophic lifestyle and its ability to manipulate its hosts sex expression (Perlin *et al*., 2015).

To survey the *S. buchloëana* genome for factors involved with host sex manipulation, we compared the colinear syntenic relationship between *S. buchloëana* and the genomic scaffolds of related *Tilletias*. Our goal was to scan for unique (non-syntenic) segments of the *S. buchloëana* genome that may have been essential to its evolution after it diverged from the *Tilletias*. The largest segment of the *S. buchloëana* genome that lacked syntenic relationship was the 106 kb short arm of acrocentric chromosome 12 that contained 44 genes, of which, 59% (26 genes) were predicted to be secreted, and six were subtilisins (Fig. 6C and E).

Chandra and Huff (2010), found that buffalograss florets show the first signs of unisexual floral development during the boot stage of inflorescences development. We compared the gene expression profiles of *S. buchloëana* grown in culture (potato dextrose agar) to *S. buchloëana* growing in the developing inflorescences of male buffalograss to identify fungal genes that might play a role in reactivating the nonfunctional pistillate rudiments of male plants. We identified 3,017 differentially expressed *S. buchloëana* genes (DEGs). Most (91%) of DEGs were downregulated *in planta* and lacked functional annotation (Supplementary Fig. 6). We mapped DEGs to the reference genome and observed that the unique short arm of chromosome 12 also contained the highest density of upregulated genes across the entire *S. buchloëana* genome (Fig. 6B and C). Of the 44 gene annotations in the short arm, 39% (17 genes) were significantly upregulated in the developing inflorescences of male plants. Although the short arm contains no syntenic relationship to the other *Tilletias*, it does have a collinear relationship with tandemly duplicated blocks across the arm (Fig. 6C and D), suggesting that sequence duplication contributed to the expansion of the short arm and may have had a major impact on the fungus’ evolution and speciation from the *Tilletia*.

We also analyzed the functional enrichments of buffalograss genes during inflorescences development and found that male plants infected with *S. buchloëana* upregulated genes involved in pistil-associated gene classes, such as nectar development, style development, and floral meristem determinacy (Fig. 6A). In addition to pistil-associated gene classes, infected buffalograss was also enriched for peptidase inhibitor activity, suggesting that *S. buchloëana* secreted peptidases (ex. serine peptidases) may have triggered some level of defense response in the host. Our analysis suggests that the short arm of chromosome 12 plays an important role in *S. buchloëana’s* host specificity and may have coevolved with buffalograss.

## Discussion

The multidimensional and extended phenotypes of biotrophic fungi and their plant hosts are complex example of parasitic manipulation of morphology. We present the chromosome-level genome assembly of *S. buchloëana,* a fungal parasite that coerces its host to develop pistils in plants that are genetically programmed not to produce such organs in order to accommodate the fungal parasite’s own reproductive biology. Our analysis suggests that *S. buchloëana* is basal to the *T. caries* and *T. walkeri* clades of fungi, having diverged *∼*40 million years ago. We find that *S. buchloëana’s* ecological novelty is likely facilitated by molecular functions encoded on the short arm of its chromosome 12, a region that is unique to *S. buchloëana,* enriched for secreted proteins and subtilisins, and has an abundance of genes that are upregulated during host floral development (Fig. 6). While some genes on the short arm of chromosome 12 may be involved with host sex manipulation or other multidimensional phenotypes, we expect that other genes are involved with biological processes that are essential for biotrophy (host penetration, defense, and evasion) in buffalograss. In addition, we identify three duplicated blocks of genes on the short arm of chromosome 12, suggesting that tandem gene duplications likely played a role in the expansion of chromosome 12 and the elevated number of subtilisins in the species. Interestingly, upon infection with *S. buchloëana*, male buffalograss upregulates genes involved in pistil development as well as peptidase inhibitors. We hypothesize that buffalograss’ upregulated pistil development genes are a *result* of manipulation, while upregulated peptidase inhibitors might be buffalograss’ defense *response* to being manipulated. Finally, we identify and characterize genetic components of the *S. buchloëana* genome, including the presence of rare inteins, biases in codon usage, and an elevated GC content. The genomic insights generated as a result of this work have led to a clearer picture of the molecular underpinnings of *S. buchloëana’s* ability to manipulate the reproductive anatomy in its plant host. This work has generated valuable genomic resources and discoveries that advances our understanding of coevolutionary dynamics and the molecular basis for disease susceptibility in cereal crops.

## Materials and Methods

### DNA extraction and Genome Sequencing

Fungal DNA was isolated from tissue grown in culture on potato dextrose agar (PDA; Alpha Biosciences Inc., Baltimore, MD) using the Fungi/Yeast Genomic DNA Isolation Kit (Norgen Biotek Corp., Ontario, Canada). High molecular weight DNA was prepared for sequencing using the SMRTbell Template Preparation kit (v.1.0), and long-read DNA sequencing was conducted using the PacBio Sequel System based on Single Molecule, Real-Time (SMRT) Sequencing technologies. The resulting BAM file was converted to FASTQ format and input into the Canu (v.1.8; Koren *et al*., 2017) *de novo* genome assembler for generation of consensus sequence and construction of pseudomolecules. The mitochondrial genome was clipped at overlapping circular ends and annotated with GeSeq (Tillich *et al*., 2017) and visualized using OGDRAW (v.1.3; Greiner *et al*., 2019) with default parameters.

### Protein Prediction and Annotation

Repetitive elements were classified using RepeatMasker (v.4.1.2) via the MAKER pipeline (Cantarel, 2008) and softmasked prior to gene annotation. Protein prediction, annotation, and genome comparisons were performed according to the Funannotate (v.1.5.1) pipeline that classifies *ab initio* gene predictions into consensus gene predictions and functionally annotates proteins. Briefly, STAR (v.2.7; Dobin *et al*., 2013) was used to align transcript evidence from the RNA-seq (see below) to the genome. Of the 18,773 initial transcript predictions, STAR aligned 7,749 to the genome. Diamond (v.0.9.22; Buchfink *et al*., 2015) and exonerate (v.2.2) were used to align UniProt’s 546,247 manually annotated and reviewed proteins (Swiss-Prot) to the *S. buchloëana* genome. Between the two tools, 1,029 preliminary alignments were identified and used for gene prediction. Transcript and protein evidence were given to the two gene predictors, GeneMark-ES (v.4.21; Brůna *et al*., 2020) and Augustus (v.3.2.1; Stanke *et al*., 2006). The resulting 12,774 gene models were passed into EvidenceModeler (v.0.1.3; Haas *et al*., 2008) and reduced to 6,555 high quality gene models. High quality models were filtered for lengths of less than 50 amino acid and the presence of transposable elements to reduce the set to 6,427 gene models. tRNAscan-SE (v.1.3.1; Lowe and Eddy, 1997) was used to identify 48 predicted tRNAs, reducing our final set of predicted genes to 6,379.

### Comparative Genomics

Fungal genomes from the Ustilaginomycotina were downloaded from NCBI and annotated in-house using the funannotate pipeline as described above to assure that downstream comparative analyses would not be biased by the annotation pipeline or other methodological restriction (See Supplementary Table 4 for the list of fungal species and isolates used). The closest fungal genus to *S. buchloëana* is the *Tilletia*. Some of the most well characterized *Tilletias* have caused economic constraints and yield loss in their cereal crop hosts (Murray and Brenan, 1998; Qin *et al*., 2021), and six of those species have publicly available reference genomes on NCBI (Castlebury, Carris, and Vanky 2005; Carris, Castebury, and Goates 2006). Briefly, *T. indica, T. caries, T. laevis,* and *T. controversa* infect wheat and have resulted in lost revenue mainly through quarantines and bans on grain imports (Nagarajan *et al*., 1997). *Tilletia walkeri* infects ryegrass species, *Lolium multiflorum* and *L. perenne* under natural conditions, and *T. horrida* causes major disease in rice and limits the use of hybrid seed production (Wang *et al,* 2018). For the functional annotation and comparative genomics of the *Tilletias, S. buchloëana*, and other Ustilaginomycotina, we queried the amino acid sequence for each set of gene annotations against the PFAM database (v.34; Bateman *et al*., 2004) to classify protein family evidence, MEROPS (v.12.3; Rawlings *et al*., 2009) for peptidases and the proteins that inhibit peptidases, the CAZyme database for families of structurally similar carbohydrate binding modules and the catalytic enzymes that alter glycosidic bonds, SignalP (v.5.0; Armenteros *et al*., 2019) to predict signaling peptides in each amino acid sequence, the COG database (v.2020; Tatusov *et al*., 2003) for clusters of orthologous genes, and antiSMASH (v.5.0; Blin *et al*., 2019) databases for functional classification of proteins. Syntenic antiSMASH clusters were visualized using the Comparative Genomics platform (CoGe; Haug-Baltzell *et al*., 2017) with the GEvo function and the LastZ algorithm for sequence alignment (Supplementary Fig. 7). Orthologous clusters between all fungal annotations were inferred using ProteinOrtho (v.6.0.16; Lechner *et al*., 2011) with parameters ‘-synteny -singles -selfblast’ to identify 328 single copy BUSCO (Simão *et al*., 2015) orthologous clusters. Subsequently, MAFFT (v.6.1; Katoh *et al*., 2013) was used to align orthologs, trimAl (v.1; Capella-Gutiérrez *et al*., 2009) to trim spurious alignments, and RAxML (v.8; Stamatakis *et al*., 2014) for phylogenetic analysis of the aligned and trimmed orthologous genes using 100 bootstraps under maximum likelihood with the flags ‘-f a -m PROTGAMMAAUTO -p 12345 -x 12345 -# 100 -n nwk’ with *Laccaria bicolor* as the specified outgroup. The resulting newick-formatted alignment file was used as input into PATHd8 (Britton *et al*., 2007) with a fixed age of the *L. bicolor* branch set to 430 Mya. PATHd8 is a rate-smoothing method that calculates substitution rates locally to scale branch lengths proportionally to the number of proposed substitutions. Intein and subtilisins were identified using annotation classes from MEROPS (Supplementary Fig. 5), aligned with MUSCLE (Edger, 2004), and phylogenetic trees were inferred by using the Maximum Likelihood method and JTT matrix-based model (Jones *et al*., 1992) in MEGAX (Kumar *et al*., 2018). Secreted proteins and effector proteins were predicted using a custom pipeline that identifies classically and non-classically secreted proteins as well as putative effector proteins. MCScanX (Wang *et al*., 2012) was used to detect colinear syntenic blocks between related *Tilletia* species and *S. buchloëana*. The collinear MCScanX file was input into SynVisio (Bandi, 2020) to visualize regions of shared homology and plot gene expression along chromosomes. The analysis of codon usage using the relative synonymous codon usage (RSCU) and the effective number of codons (ENc) were calculated on an orthologous *Tilletia* gene set (that included at least 948 of 985 orthologous genes depending on the species) using CAIcal (Puigbò *et al*., 2008) to calculate the codon adapter index and measure the synonymous codon usage bias for orthologous pairs of genes. The predicted theoretical ENc was calculated using GC3 values and the formula, 2+GC3+29/((GC3*GC3)+((1-GC3)*(1-GC3))).We also used CodonW for multivariate comparisons of codon and amino acid usage with default parameters. Repeat-induced point mutations in the *S. buchloëana* genome were identified using RIPper (van Wyk *et al*., 2019).

### Centromere Annotation

Mauve (Darling *et al*., 2004) was used to compare sequence identity to genomic scaffolds of *Tilletia* species using a multiple genome alignment. Gene density and GC content were plotted along a 25 kb and 500 bp sliding window, respectively. Only ten of the 22 *S. buchloëana* chromosomes showed a marked and sustained decrease in GC content, generally approaching a level of 54% GC content somewhere along the chromosome, indicating a potential location for a centromeric region. Chromosomes with a visible drop in GC content (ex. chromosomes 2, 9, 11, and 15) also displayed the lowest overall gene density within the same stretch of chromosome as well as distinct clusters in terms of length and number of LTRs (Fig. 2; Supplementary Fig. 3). With four exceptions, we found that the length of LTRs gave the best definition of the centromeric boundaries such that the first LTR greater than 388 bp in length (*i.e.*, the shortest LTR in the candidate centromeric region) from either end of the chromosome marked the centromeric beginning/end. In addition, several of the chromosomes showed two adjacent clusters of LTRs greater than 500 bp within the centromeric region along with an associated decrease of gene density (ex. chromosomes 3, 6, and 10) indicting a possible cluster of LTRs on either side of the actual centromere. Taken together, the convergence of low GC content, low gene density, and the high frequency of LTRs (primarily Copia and Gypsy; Supplementary Table 2) greater than 388 bp were used to predict the locations of *S. buchloëana’s* centromeric regions (Fig. 2; Supplementary Fig. 3). Among the four exceptions to this centromeric boundary ‘rule’, two involved rDNA located in the telocentric regions of chromosomes 3 and 22 that contained several LTRs greater that 388 bp, one involved only a single LTR on chromosome 6 which was of the Bel/Pao family, and one involved a cluster of numerous long LTRs at the telocentric region of chromosome 10 and may represent a new LTR invasion.

### RNA-seq

A transcriptomic RNA-seq analysis was performed comparing a population of 28 male buffalograss genotypes that were either infected or mock-infected with *S. buchloëana* teliospores. The genotypes utilized were the same 28 male genotypes evaluated in a previous study (Chandra and Huff, 2014). Immature (boot-stage) inflorescences, approximately 3 to 7 mm in length, were harvested from either infected or mock-infected plants in the afternoon (3-5 pm) every day for approximately three weeks and immediately placed in liquid nitrogen and stored at -80 C. After tissues were harvested, treatment combinations were pooled, lyophilized, and stored at -20 C for approximately six years. RNA was extracted from four biological replicates with tissue samples from each treatment for a total of eight RNA samples. RNA extractions were verified for adequate quality and concentration using a Bioanalyzer (Agilent Technologies, California, USA). Samples with an RNA Integrity Number (RIN) of 6.8 or higher were sent to the Pennsylvania State Genomics Core Facility for sequencing using an Illumina MiSeq and 150×150 bp pair-end libraries.

Sequences were trimmed for adapters and low-quality ends using bbduk with parameters ‘tbo tpe ktrim = r k = 23 mink = 11 hdist = 1’. Cleaned sequences from uninfected buffalograss and *S. buchloëana* grown in culture were input into the trinityrnaseq toolkit (v2.13.0; Haas *et al*., 2013) to assemble *de novo* transcriptomes. Reads from infected plants contained both buffalograss and *S. buchloëana* sequences, and so they were subsequently aligned to the reference transcriptomes of both species to separate transcripts based on their species of origin (Supplementary Fig. 8). Kallisto (Bray *et al*., 2016) and DESeq2 (Love *et al*., 2014) were then executed using the trinityrnaseq scripts ‘align_and_estimate_abundance.pl’ and ‘run_DE_analysis.pl’ to conduct the differential expression analysis. Functional annotations were assigned using Trinotate (trinityrnaseq toolkit) and the id2go formatted file was analyzed using the ‘analyze_diff_expr.pl’ with the ‘–examine_GO_enrichment’ flag to call Goseq to examine functionally enriched gene ontologies. Gene ontologies were visualized using Revigo (Supek *et al*., 2011) to cluster enriched ontologies by semantic similarity.

## Data Availability Statement

Genome assembly and gene annotation files are publicly available through the CyVerse CoGe platform (https://genomevolution.org/coge/). Raw sequence data are available in the Sequence Read Archive under NCBI BioProject PRJNA961724.

## Acknowledgements

None.

## Literature Cited

1. Almagro Armenteros JJ et al. 2019. SignalP 5.0 improves signal peptide predictions using deep neural networks. Nat Biotechnol. 37:420–423.

2. Bandi VK. 2020. SynVisio: A Multiscale Tool to Explore Genomic Conservation. Thesis, University of Saskatchewan.

3. Bateman A et al. 2004. The Pfam protein families database. Nucleic Acids Res. 32:D138–D141.

4. Blin K et al. 2019. antiSMASH 5.0: updates to the secondary metabolite genome mining pipeline. Nucleic Acids Res. 47:W81–W87.

5. Bray NL, Pimentel H, Melsted P, Pachter L. 2016. Near-optimal probabilistic RNA-seq quantification. Nat Biotechnol. 34:525–527.

6. Britton T, Anderson CL, Jacquet D, Lundqvist S, Bremer K. 2007. Estimating Divergence Times in Large Phylogenetic Trees. Systematic Biology. 56:741–752.

7. Brůna T, Hoff KJ, Lomsadze A, Stanke M, Borodovsky M. 2021. BRAKER2: automatic eukaryotic genome annotation with GeneMark-EP+ and AUGUSTUS supported by a protein database. NAR Genomics and Bioinformatics. 3:lqaa108.

8. Buchfink B, Xie C, Huson DH. 2015. Fast and sensitive protein alignment using DIAMOND. Nat Methods. 12:59–60.

9. Cantarel BL et al. 2008. MAKER: an easy-to-use annotation pipeline designed for emerging model organism genomes. Genome Res. 18:188–196.

10. Capella-Gutiérrez S, Silla-Martínez JM, Gabaldón T. 2009. trimAl: a tool for automated alignment trimming in large-scale phylogenetic analyses. Bioinformatics. 25:1972–1973.

11. Carris LM, Castlebury LA, Goates BJ. 2006. Nonsystemic Bunt Fungi—*Tilletia* indica and *T. horrida*: A Review of History, Systematics, and Biology. Annual Review of Phytopathology. 44:113–133.

12. Castlebury LA, Carris LM, Vánky K. 2005. Phylogenetic analysis of *Tilletia* and allied genera in order Tilletiales (Ustilaginomycetes; Exobasidiomycetidae) based on large subunit nuclear rDNA sequences. Mycologia. 97:888–900.

13. Cézilly F, Favrat A, Perrot-Minnot M-J. 2013. Multidimensionality in parasite-induced phenotypic alterations: ultimate versus proximate aspects. Journal of Experimental Biology. 216:27–35.

14. Chandra A, Huff DR. 2008. *Salmacisia*, a New Genus of Tilletiales: Reclassification of *Tilletia buchloëana* Causing Induced Hermaphroditism in Buffalograss. Mycologia. 100:81–93.

15. Chandra A, Huff DR. 2010. A Fungal Parasite Regulates a Putative Female-Suppressor Gene Homologous to Maize *Tasselseed2* and Causes Induced Hermaphroditism in Male Buffalograss. MPMI. 23:239–250.

16. Chandra A, Huff DR. 2014. Pistil Smut Infection Increases Ovary Production, Seed Yield Components, and Pseudosexual Reproductive Allocation in Buffalograss. Plants. 3:594– 612.

17. Darling ACE, Mau B, Blattner FR, Perna NT. 2004. Mauve: Multiple Alignment of Conserved Genomic Sequence With Rearrangements. Genome Res. 14:1394–1403.

18. Dawkins R. 2016. The Extended Phenotype: The Long Reach of the Gene. Oxford University Press.

19. Diner RE et al. 2017. Diatom centromeres suggest a mechanism for nuclear DNA acquisition. PNAS. 114:E6015–E6024.

20. Dobin A et al. 2013. STAR: ultrafast universal RNA-seq aligner. Bioinformatics. 29:15– 21.

21. Duan X, Gimble FS, Quiocho FA. 1997. Crystal Structure of PI-SceI, a Homing Endonuclease with Protein Splicing Activity. Cell. 89:555–564.

22. Edgar RC. 2004. MUSCLE: multiple sequence alignment with high accuracy and high throughput. Nucleic Acids Research. 32:1792–1797.

23. Felsenstein J. 1978. Cases in which Parsimony or Compatibility Methods will be Positively Misleading. Systematic Biology. 27:401–410.

24. Fryxell KJ, Zuckerkandl E. 2000. Cytosine Deamination Plays a Primary Role in the Evolution of Mammalian Isochores. Molecular Biology and Evolution. 17:1371–1383.

25. Green CM, Novikova O, Belfort M. 2018. The dynamic intein landscape of eukaryotes. Mobile DNA. 9:4.

26. Greiner S, Lehwark P, Bock R. 2019. OrganellarGenomeDRAW (OGDRAW) version 1.3.1: expanded toolkit for the graphical visualization of organellar genomes. Nucleic Acids Research. 47:W59–W64.

27. Haas BJ et al. 2008. Automated eukaryotic gene structure annotation using EVidenceModeler and the Program to Assemble Spliced Alignments. Genome Biology. 9:R7.

28. Haas BJ et al. 2013. De novo transcript sequence reconstruction from RNA-seq using the Trinity platform for reference generation and analysis. Nat Protoc. 8:1494–1512.

29. Hane JK, Oliver RP. 2008. RIPCAL: a tool for alignment-based analysis of repeat-induced point mutations in fungal genomic sequences. BMC Bioinformatics. 9:478.

30. Haug-Baltzell A, Stephens SA, Davey S, Scheidegger CE, Lyons E. 2017. SynMap2 and SynMap3D: web-based whole-genome synteny browsers. Bioinformatics. 33:2197–2198.

31. He M-Q et al. 2019. Notes, outline and divergence times of Basidiomycota. Fungal Diversity. 99:105–367.

32. Henry LP, Bruijning M, Forsberg SKG, Ayroles JF. 2021. The microbiome extends host evolutionary potential. Nat Commun. 12:5141.

33. Hood ME et al. 2010. Distribution of the anther-smut pathogen *Microbotryum* on species of the Caryophyllaceae. New Phytol. 187:217–229.

34. Huff DR, Zagory D, and Wu L. 1987. Report of buffalograss bunt found in Oklahoma. Plant Disease. 71:651.

35. Jayawardena RS et al. 2019. One stop shop II: taxonomic update with molecular phylogeny for important phytopathogenic genera: 26–50 (2019). Fungal Diversity. 94:41–129.

36. Jiang RHY et al. 2013. Distinctive Expansion of Potential Virulence Genes in the Genome of the Oomycete Fish Pathogen *Saprolegnia parasitica*. PLOS Genetics. 9:e1003272.

37. Jones DT, Taylor WR, Thornton JM. 1992. The rapid generation of mutation data matrices from protein sequences. Comput Appl Biosci. 8:275–282.

38. Kämper J et al. 2006. Insights from the genome of the biotrophic fungal plant pathogen *Ustilago maydis*. Nature. 444:97–101.

39. Katoh K, Standley DM. 2013. MAFFT Multiple Sequence Alignment Software Version 7: Improvements in Performance and Usability. Molecular Biology and Evolution. 30:772–780.

40. Kellerman WA, Swingle WT. 1889. New Species of Kansas Fungi. The Journal of Mycology. 5:11–49.

41. Kemler M et al. 2020. Host preference and sorus location correlate with parasite phylogeny in the smut fungal genus *Microbotryum* (Basidiomycota, Microbotryales). Mycol Progress. 19:481–493.

42. Kinney M, Columbus T, Friar E. 2007. Dicliny in *Bouteloua* (Poaceae: Chloridoideae): Implications for the Evolution of Dioecy. Aliso. 23:605–614.

43. Koren S et al. 2017. Canu: scalable and accurate long-read assembly via adaptive k-mer weighting and repeat separation. Genome Res. 27:722–736.

44. Kumar S, Stecher G, Li M, Knyaz C, Tamura K. 2018. MEGA X: Molecular Evolutionary Genetics Analysis across Computing Platforms. Mol Biol Evol. 35:1547– 1549.

45. Lechner M et al. 2011. Proteinortho: Detection of (Co-)orthologs in large-scale analysis. BMC Bioinformatics. 12:124.

46. Leger RJS, Joshi L, Roberts DW. 1997. Adaptation of proteases and carbohydrases of saprophytic, phytopathogenic and entomopathogenic fungi to the requirements of their ecological niches. Microbiology. 143:1983–1992.

47. Li X-Q, Du D. 2014. Variation, Evolution, and Correlation Analysis of C+G Content and Genome or Chromosome Size in Different Kingdoms and Phyla. PLOS ONE. 9:e88339.

48. Long H et al. 2018. Evolutionary determinants of genome-wide nucleotide composition. Nat Ecol Evol. 2:237–240.

49. Love MI, Huber W, Anders S. 2014. Moderated estimation of fold change and dispersion for RNA-seq data with DESeq2. Genome Biology. 15:550.

50. Lowe TM, Eddy SR. 1997. tRNAscan-SE: A Program for Improved Detection of Transfer RNA Genes in Genomic Sequence. Nucleic Acids Research. 25:955–964.

51. Melters DP et al. 2013. Comparative analysis of tandem repeats from hundreds of species reveals unique insights into centromere evolution. Genome Biology. 14:R10.

52. Meyne J et al. 1990. Distribution of non-telomeric sites of the (TTAGGG)n telomeric sequence in vertebrate chromosomes. Chromosoma. 99:3–10.

53. Murray GM, Brenan JP. 1998. The risk to Australia from *Tilletia indica*, the cause of Karnal bunt of wheat. Australasian Plant Pathology. 27:212–225.

54. Muszewska A et al. 2017. Fungal lifestyle reflected in serine protease repertoire. Sci Rep. 7:1–12.

55. Nagarajan S et al. 1997. Karnal bunt (*Tilletia indica*) of wheat – a review. Review of Plant Pathology. 76:8.

56. Park BH, Karpinets TV, Syed MH, Leuze MR, Uberbacher EC. 2010. CAZymes Analysis Toolkit (CAT): Web service for searching and analyzing carbohydrate-active enzymes in a newly sequenced organism using CAZy database. Glycobiology. 20:1574– 1584.

57. Perlin MH et al. 2015. Sex and parasites: genomic and transcriptomic analysis of *Microbotryum lychnidis-dioicae*, the biotrophic and plant-castrating anther smut fungus. BMC Genomics. 16:461.

58. Piątek M, Riess K, Karasiński D, Yorou NS, Lutz M. 2016. Integrative analysis of the West African *Ceraceosorus africanus* sp*. nov.* provides insights into the diversity, biogeography, and evolution of the enigmatic Ceraceosorales (Fungi: Ustilaginomycotina). Org Divers Evol. 16:743–760.

59. Poulin R. 2013. Parasite manipulation of host personality and behavioural syndromes. Journal of Experimental Biology. 216:18–26.

60. Puigbò P, Bravo IG, Garcia-Vallve S. 2008. CAIcal: A combined set of tools to assess codon usage adaptation. Biol Direct. 3:38.

61. Qin DD, Xu TS, Liu TG, Chen WQ, Gao L. 2021. First Report of Wheat Common Bunt Caused by *Tilletia laevis* in Henan Province, China. Plant Disease. 105:215.

62. Rawlings ND et al. 2018. The MEROPS database of proteolytic enzymes, their substrates and inhibitors in 2017 and a comparison with peptidases in the PANTHER database. Nucleic Acids Res. 46:D624–D632.

63. Selker EU, Stevens JN. 1985. DNA methylation at asymmetric sites is associated with numerous transition mutations. Proceedings of the National Academy of Sciences. 82:8114–8118.

64. Shah NH, Muir TW. 2014. Inteins: Nature’s Gift to Protein Chemists. Chem Sci. 5:446– 461.

65. Sharma R, Xia X, Riess K, Bauer R, Thines M. 2015. Comparative Genomics Including the Early-Diverging Smut Fungus *Ceraceosorus bombacis* Reveals Signatures of Parallel Evolution within Plant and Animal Pathogens of Fungi and Oomycetes. Genome Biology and Evolution. 7:2781.

66. Simão FA, Waterhouse RM, Ioannidis P, Kriventseva EV, Zdobnov EM. 2015. BUSCO: assessing genome assembly and annotation completeness with single-copy orthologs. Bioinformatics. 31:3210–3212.

67. Smith MM. 2002. Centromeres and variant histones: what, where, when and why? Curr. Opin. Cell Biol. 14:279–285.

68. Stamatakis A. 2014. RAxML version 8: a tool for phylogenetic analysis and post-analysis of large phylogenies. Bioinformatics. 30:1312–1313.

69. Stanke M et al. 2006. AUGUSTUS: *ab initio* prediction of alternative transcripts. Nucleic Acids Research. 34:W435–W439.

70. Storck R. 1966. Nucleotide composition of nucleic acids of fungi. II. Deoxyribonucleic acids. J Bacteriol. 91:227–230.

71. Sun G, Pourkheirandish M, Komatsuda T. 2009. Molecular evolution and phylogeny of the RPB2 gene in the genus Hordeum. Ann Bot. 103:975–983.

72. Supek F, Bošnjak M, Škunca N, Šmuc T. 2011. REVIGO summarizes and visualizes long lists of gene ontology terms. PLoS One. 6:e21800.

73. Tatusov RL et al. 2003. The COG database: an updated version includes eukaryotes. BMC Bioinformatics. 4:41.

74. Thomas F, Poulin R, Brodeur J. 2010. Host manipulation by parasites: a multidimensional phenomenon. Oikos. 119:1217–1223.

75. Tillich M et al. 2017. GeSeq – versatile and accurate annotation of organelle genomes. Nucleic Acids Research. 45:W6–W11.

76. Uchida W, Matsunaga S, Sugiyama R, Kazama Y, Kawano S. 2003. Morphological development of anthers induced by the dimorphic smut fungus *Microbotryum violaceum* in female flowers of the dioecious plant *Silene latifolia*. Planta. 218:240–248.

77. van Houte S, Ros VID, van Oers MM. 2013. Walking with insects: molecular mechanisms behind parasitic manipulation of host behaviour. Molecular Ecology. 22:3458–3475.

78. van Wyk S et al. 2019. The RIPper, a web-based tool for genome-wide quantification of Repeat-Induced Point (RIP) mutations. PeerJ. 7:e7447.

79. Vinogradov AE. 2003. DNA helix: the importance of being GC-rich. Nucleic Acids Res. 31:1838–1844.

80. Vyas A. 2015. Mechanisms of Host Behavioral Change in *Toxoplasma gondii* Rodent Association. PLOS Pathogens. 11:e1004935.

81. Wang A et al. 2018. The pathogenic mechanisms of Tilletia horrida as revealed by comparative and functional genomics. Scientific Reports. 8.

82. Wang Y et al. 2012. MCScanX: a toolkit for detection and evolutionary analysis of gene synteny and collinearity. Nucleic Acids Research. 40:e49–e49.

83. Werren JH. 2011. Selfish genetic elements, genetic conflict, and evolutionary innovation. Proceedings of the National Academy of Sciences. 108:10863–10870.

84. Wolfe KH, Sharp PM, Li WH. 1989. Mutation rates differ among regions of the mammalian genome. Nature. 337:283–285.

85. Wu P, van Overbeek M, Rooney S, de Lange T. 2010. Apollo Contributes to G Overhang Maintenance and Protects Leading-End Telomeres. Molecular Cell. 39:606–617.

86. Zhao R-L et al. 2017. A six-gene phylogenetic overview of Basidiomycota and allied phyla with estimated divergence times of higher taxa and a phyloproteomics perspective. Fungal Diversity. 84:43–74.

